# Locomotion-induced neural activity independent of auditory feedback in the mouse inferior colliculus

**DOI:** 10.1101/2025.05.27.656503

**Authors:** Jisoo Han, Haiyan Jiang, Young Rae Ji, Gunsoo Kim

**Affiliations:** Sensory and motor systems Research Group, Korea Brain Research Institute (KBRI), Daegu, 41068, South Korea; Center for Neuroscience Imaging (CNIR), Institute for Basic Science (IBS), Suwon, 16419, South Korea; Department of Biomedical Engineering, Sungkyunkwan University, Suwon, 16419, South Korea

## Abstract

Accumulating evidence indicates that the auditory pathway integrates movement-related signals with auditory input, yet the precise sources and mechanisms of this integration across various processing levels are incompletely understood. The inferior colliculus (IC), a major midbrain hub in the auditory pathway, shows widespread modulation of neural activity during locomotion, indicating that auditory neurons at this level are sensitive to ongoing movement. However, in hearing animals, it has been challenging to dissociate auditory feedback from other motor-related signals. In this study, to isolate non-auditory contributions, we recorded IC neural activity during locomotion in deafened mice, thereby eliminating all auditory feedback through both air and bone conduction. Even in the absence of auditory input, IC neurons exhibited robust, bidirectional modulation during locomotion. Timing analysis using electromyography revealed both predictive and feedback components relative to locomotion onset. Furthermore, the timing and direction of modulation varied considerably across different locomotion bouts, suggesting convergence of multiple non-auditory inputs. These findings demonstrate that non-auditory, movement-related signals significantly shape auditory midbrain activity through both predictive and feedback mechanisms.

## INTRODUCTION

Locomotion is a fundamental behavior across species and strongly influences neural activity of the auditory pathway. Prior studies have shown that both spontaneous and sound-evoked activity are modulated during locomotion [1–10]. During navigation, for example, this modulation may serve important functions, such as encoding movement speed [10,11] and distinguishing self-generated sounds from those in the environment [12–14]. While locomotion-related modulation has been well characterized in the auditory cortex [1–3,5,10], similar modulation has also been observed at subcortical levels [4,6,7]. However, how these signals are distributed across different processing stages of the auditory pathway remains poorly understood.

The inferior colliculus (IC) serves as a major midbrain hub that integrates auditory and other multisensory inputs [15,16]. Recent studies have shown that neural activity in the IC is robustly modulated by locomotion, indicating that midbrain auditory neurons are sensitive to ongoing movement [6,7]. Significant modulation was observed in approximately 75% of IC neurons, with two thirds showing increased firing and one third showing suppression. These modulations occur not only in multisensory regions such as lateral and dorsal cortices (LCIC and DCIC), but also in the central nucleus (CNIC), traditionally considered a primarily auditory region [17–21]. While these findings suggest that processing movement-related information is a fundamental function of the IC, the sources and specific roles of this modulation remain unclear.

To understand the contribution of specific inputs, it is essential to isolate the effects of auditory and non-auditory movement-related signals. Locomotion generates sounds that provide auditory feedback, which can influence sound processing or ongoing movement itself [13,22]. In the IC, such feedback likely contributes to the observed modulation, especially in the CNIC. However, prior studies in hearing mice could not separate the effects of auditory feedback from other motor-related signals, making it difficult to identify the role of non-auditory inputs.

While locomotion-related modulation has been reported in hearing-impaired mice [7], residual auditory function due to incomplete deafening complicates interpretation. Moreover, movement can induce skull vibrations that stimulate cochlear hair cells via bone conduction [23], providing an additional pathway for auditory feedback. The potential contribution of this mechanism has not yet been examined.

In this study, we recorded IC neural activity during locomotion in mice that can be rendered deaf through diphtheria toxin (DT) treatment (Pou4f3^+/DTR^ line;[24]), thereby eliminating auditory feedback through both air and bone conduction. We observed clear, bidirectional modulation of IC activity during locomotion, even in the absence of auditory input. Timing analysis using electromyography revealed that these modulations include both anticipatory and feedback components. Furthermore, substantial variability in modulation patterns across locomotion bouts suggests convergence of diverse non-auditory signals. Our findings demonstrate that non-auditory, movement-related signals significantly shape neural activity in the auditory midbrain, engaging both predictive and feedback mechanisms.

## RESULT

### Locomotion-induced auditory feedback in head-fixed mice

We first investigated whether locomotion in head-fixed mice on a passive treadmill generates significant auditory feedback. To assess air-conducted auditory feedback, we recorded locomotion-generated sounds using a microphone positioned near the mouse’s head (Fig 1A). Soft footstep sounds were detected during locomotion bouts, with most of the sound energy concentrated between 2 and 10 kHz (Fig 1B and 1C). The average sound intensity of individual footsteps was 40 ± 3dB SPL (109 events; range: 35∼51dB). To assess whether locomotion induces significant vibrations under head-fixed conditions, we also measured skull vibrations using a vibration sensor attached to the skull (Fig 1D; [25]). Clear vibration signals were detected during walking bouts and were strongly correlated with walking speed (Fig 1E and 1F; r = 0.43 ± 0.09, N = 3 mice). These results demonstrate that, even under head-fixed conditions, locomotion generates substantial auditory feedback through both airborne and bone-conducted pathways, likely contributing to neural modulation in the auditory pathway during movement.

**Fig 1.**
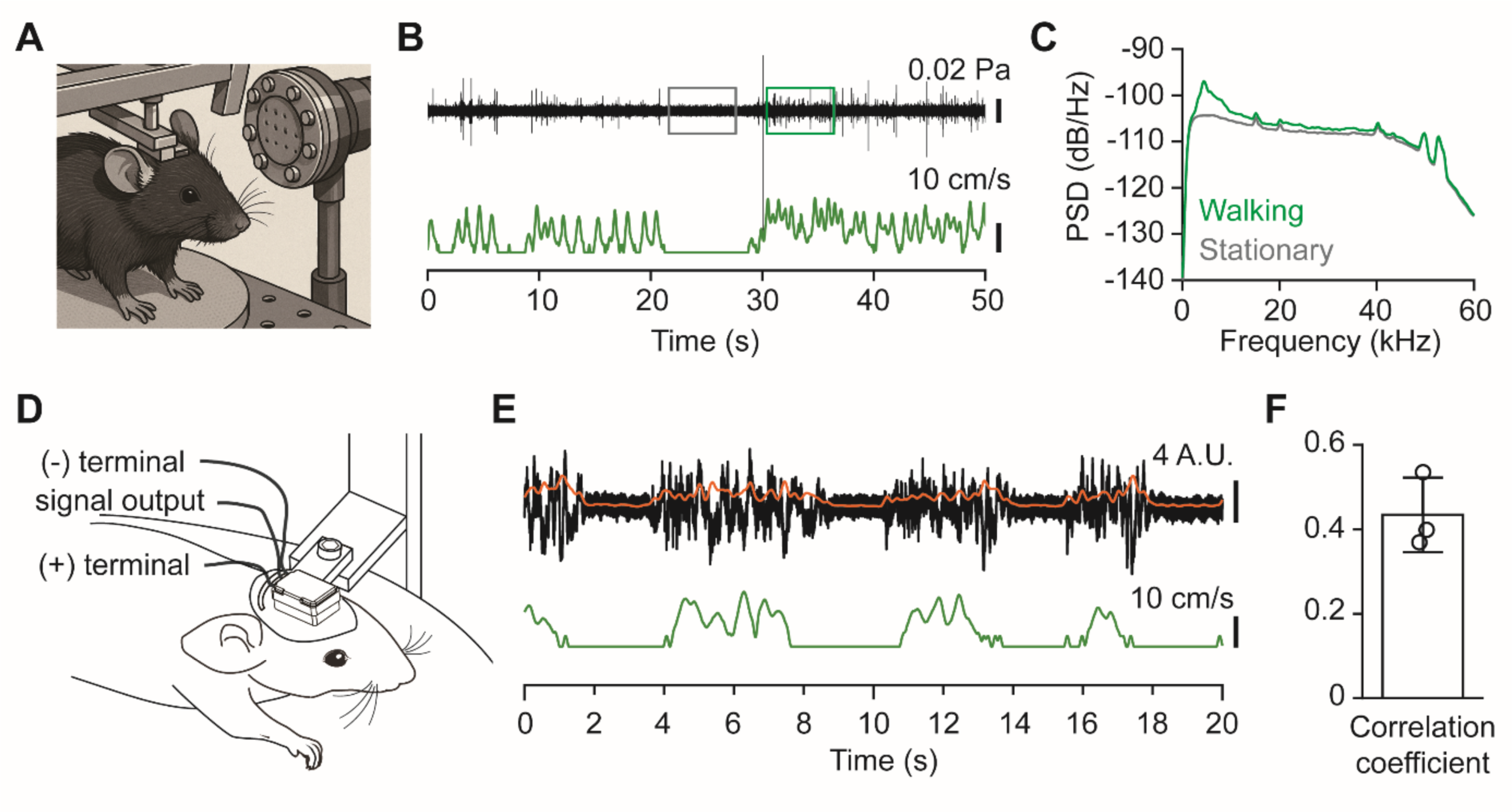
Locomotion generates detectable auditory feedback in head-fixed mice. (**A**) Experimental setup for recording footstep-generated sounds in a head-fixed mouse on a passive treadmill. (**B**) Top: sound pressure waveform during walking (black trace). Boxes indicate stationary (gray) and walking (green) periods used for analysis in (C). Bottom: corresponding walking speed (green trace). (**C**) Power spectral density of sound recordings during stationary (gray) and walking (green) periods. (**D**) Schematic of the skull-mounted sensor setup for detecting bone-conducted vibrations in head-fixed mice during walking. (**E**) Example traces of vibration sensor signal (black, bandpass; orange, smoothed) and walking speed (green). (**F**) Correlation between smoothed sensor signal and walking speed. Bar graph shows the mean correlation coefficient ± SD (N = 3 mice).

### Locomotion modulates IC neural activity in the absence of auditory feedback

To eliminate auditory feedback during locomotion and isolate non-auditory components of neural modulation in the auditory pathway, we used Pou4f3^+/DTR^ mice, which express the diphtheria toxin receptor (DTR) in cochlear hair cells [24]. One week after diphtheria toxin (DT) injection (25 ng/Kg), we assessed deafness using whole-mount cochlear histology and neural recordings. Myosin VIIa immunostaining revealed a near-complete loss of hair cells throughout all cochlear turns in DTR mice (Fig 2A). Consistent with this anatomical result, multi-unit recordings from the IC showed no sound-evoked responses (Fig 2B, 2C and S1 Fig). The absence of cochlear hair cells should also prevent locomotion-induced skull vibrations from being transduced into neural signals via bone conduction [23,26]. These results confirm that DT injection in Pou4f3^+/DTR^ mice effectively induces deafness (hereafter referred to as “deaf mice”), eliminating auditory feedback during locomotion.

**Fig 2.**
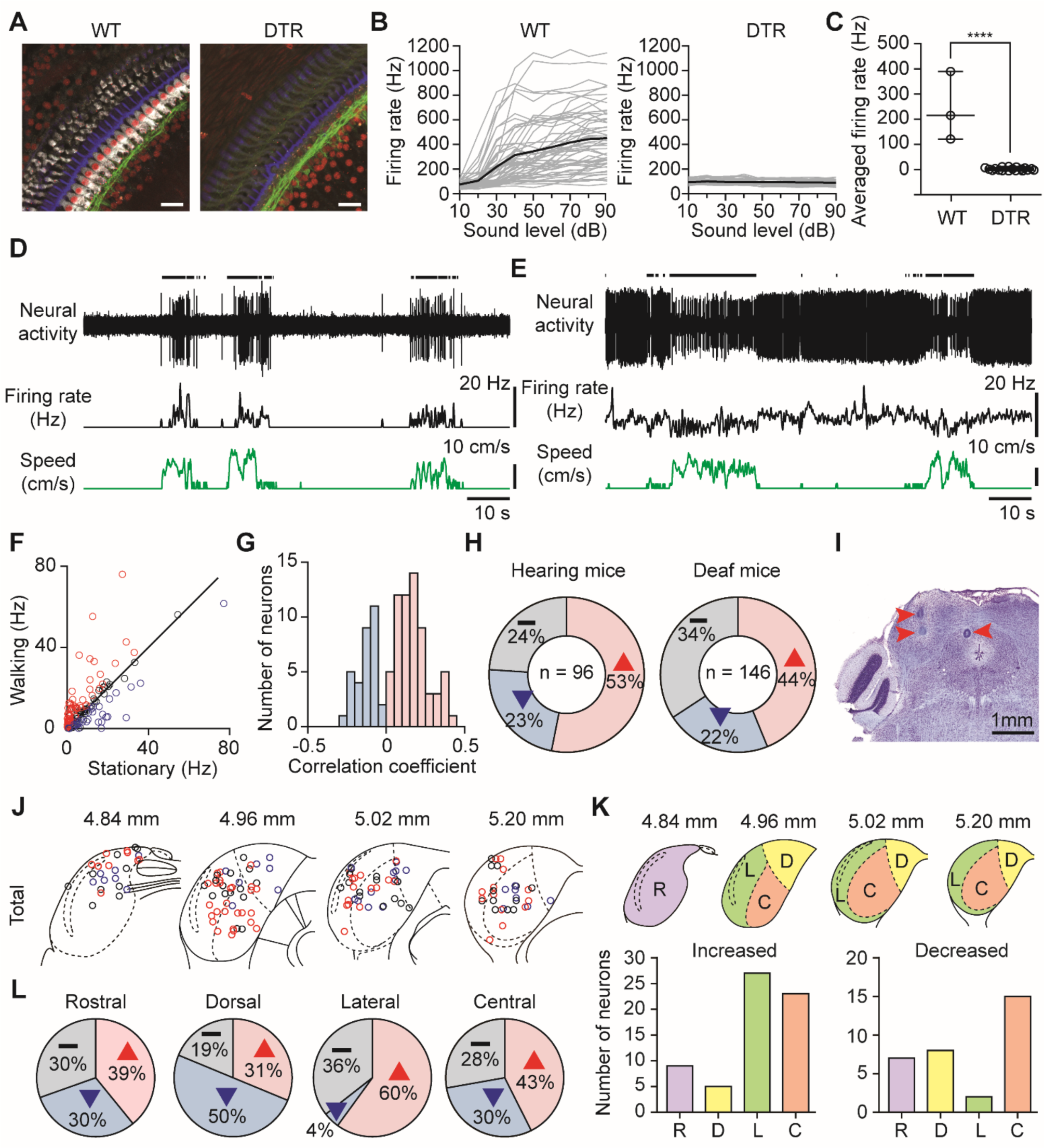
Modulation of IC neural activity during locomotion in deaf mice. (**A**) Representative images of the cochlear apical turn from a WT mouse (left) and the middle turn from a DTR mouse (right). In DTR mice injected with diphtheria toxin, hair cells (labeled with myosin VIIa, gray) are absent, confirming successful ablation. Additional markers: ribbon synapses (CtBP2, red), afferent nerve fibers (Tuj1, green), and hair cell stereocilia (phalloidin-labeled F-actin, blue). Scale bar: 20 µm. (**B**) Sound-evoked IC multi-unit firing rates across sound levels (dB) in a WT (left) and a DTR (right) mice. Each gray trace represents responses from a single recording site; thick black line indicates the average. (**C**) Comparison of average firing rates at 70 dB SPL in WT and DTR mice. Median value of average firing rates are 215.4 Hz (WT) and 1.6 Hz (DTR) mice (WT: N = 3 mice; DTR: N = 18 mice; **** p < 0.0001, Welch’s t-test). (**D** and **E**) Example IC single-unit recordings in deaf mice exhibiting increased (D) or decreased (E) firing rates during locomotion. Top: raw spike trace; middle: smoothed firing rate; bottom: walking speed. Black bars (above top trace) indicate walking periods. (**F**) Scatter plot of mean firing rates during stationary vs. walking periods for all IC neurons recorded in deaf mice (n = 146 neurons). Red: increased firing; blue: decreased firing; black: no modulation. Diagonal line indicates unity. (**G**) Distribution of correlation coefficients between firing rate and walking speed for modulated neurons. Pink bars: increased firing; blue bars: decreased firing. (**H**) Proportions of neurons in hearing (left, from[7]) and deaf (right) mice with increased (pink), decreased (blue), or unchanged (gray) activity during locomotion. (**I**) Nissl-stained coronal section showing lesion sites (red arrowheads) used to verify IC recording locations. Scale bar: 1 mm. (**J**) Spatial distribution of recorded IC neurons across four coronal sections (4.84, 4.96, 5.02, 5.20 mm posterior to bregma). Red: increased firing; blue: decreased firing; black: unmodulated neurons. (**K**) IC subdivisions – rostral (R), dorsal (D), lateral (L), and central (C) – were classified in different colors. Proportion of increased (left) and decreased (right) locomotion-modulated neurons across IC subdivisions. (**L**) Proportion of modulation types within each subdivision. Pink: increase firing; blue: decreased; gray: no modulation.

To investigate whether IC neural activity is modulated by locomotion in the absence of auditory feedback, we performed single-unit recordings in head-fixed, deaf mice on a passive treadmill. Deafness was confirmed in each animal by the absence of sound-evoked responses to broadband noise stimuli presented during the experiment (Fig 2C).

Deafening significantly decreased baseline activity (S2A Fig), but we observed clear locomotion-induced modulation in IC neural activity. In an example IC neuron, the average firing rate increased from 0.2 Hz at rest to 3.8 Hz during locomotion (Fig 2D). In contrast, another neuron showed a decrease from 10.5 Hz to 8.0 Hz during locomotion (Fig 2E). The average change in firing rate was comparable between hearing and deaf mice (S2B Fig; Increase: 14.0 Hz vs 7.5 Hz; Decrease: −7.9 Hz vs −6.8 Hz), resulting in greater percent modulation in both excited and suppressed IC neurons in deaf mice (S2C Fig; Increase: 257% vs 537%; Decrease: −39% vs −54%).

Among the 146 IC neurons recorded, 64 neurons (44%) increased their firing during locomotion, exhibiting positive correlations with walking speed, whereas 32 neurons (22%) decreased their firing, exhibiting negative correlations with speed (Fig 2F, 2G and 2H). Compared to hearing mice, the proportion of neurons with excitatory modulation decreased from 53% to 44% (∼20% reduction), accompanied by an increase of unmodulated neurons (from 24% to 34%; Fig 2H). This shift likely reflects the absence of auditory feedback, including bone conduction. The portion of suppressed neurons did not change, suggesting that auditory feedback has little effect on locomotion-induced suppression of IC activity. These results demonstrate that IC neural activity is robustly modulated by locomotion, even in the absence of auditory feedback, establishing that the majority of this modulation originates from non-auditory sources.

The shell regions of the IC (LCIC and DCIC) are known to be multisensory, receiving somatosensory inputs, for example, in addition to the auditory inputs [20,21]. Therefore, one might expect that non-auditory component of locomotion-induced activity modulation occurs primarily in the shell regions. We investigated the spatial distribution of neurons modulated by locomotion across the IC by dividing it into four subdivisions in deaf mice: rostral (R), lateral (L), dorsal (D), and central (C) (Fig 2J; [27]). Our reconstruction of recording locations based on lesions (Fig 2I) revealed that locomotion-induced modulation is observed in all IC subdivisions in deaf mice. While most modulated IC neurons were located in the shell region, a significant portion was also found in the CNIC for both directions of modulation (64% (shell) vs 36% (core) for the increased; 53% (shell) vs 47% (core) for the decreased; Fig 2K).

We also examined the proportions of modulated neurons and their modulation directions in each subregion. In the LCIC, excitatory modulation was predominant, while suppressive modulation was rare. Conversely, in the DCIC, suppressive modulation was most prominent. Across all subdivisions, more than 60% of the neurons exhibited sensitivity to locomotion (Fig 2L). These findings demonstrate that locomotion-induced activity modulation occurs across all IC subdivisions, even in deafness, with regional differences in prevalence of the direction of modulation.

### Temporal relationship between IC neural activity modulation and locomotion onset

Neural modulation that precedes movement onset is consistent with a predictive signal such as corollary discharge [1], whereas modulation that follows movement onset is more consistent with a feedback signal such as somatosensory inputs induced by movement [28,29]. Previous studies have shown that in many IC neurons, neural modulation precedes locomotion onset. Here, we asked whether the absence of auditory feedback in deaf mice influences the timing of modulation.

We determined modulation onset times for each neuron by averaging firing rates across individual walking bouts, as shown in the example neurons (Fig 3A, 3B and 3C). In neurons with increased firing, modulation began significantly earlier in deaf mice compared to hearing mice (Fig 3D; median latency: −179 ms (deaf) vs −110 ms (hearing)). In contrast, for neurons with decreased firing during locomotion, the timing of modulation onset was similar between the two groups (Fig 3D; median latency: 40 ms (deaf) vs 26 ms (hearing)). In both deaf and hearing mice, excitatory modulation was more likely to begin before locomotion onset, whereas suppressive modulation tended to occur either slightly before or after movement began (Fig 3E). We observed no clear patterns in modulation latency across IC subregions (S3 Fig), suggesting that timing characteristics are broadly distributed throughout the IC.

**Fig 3.**
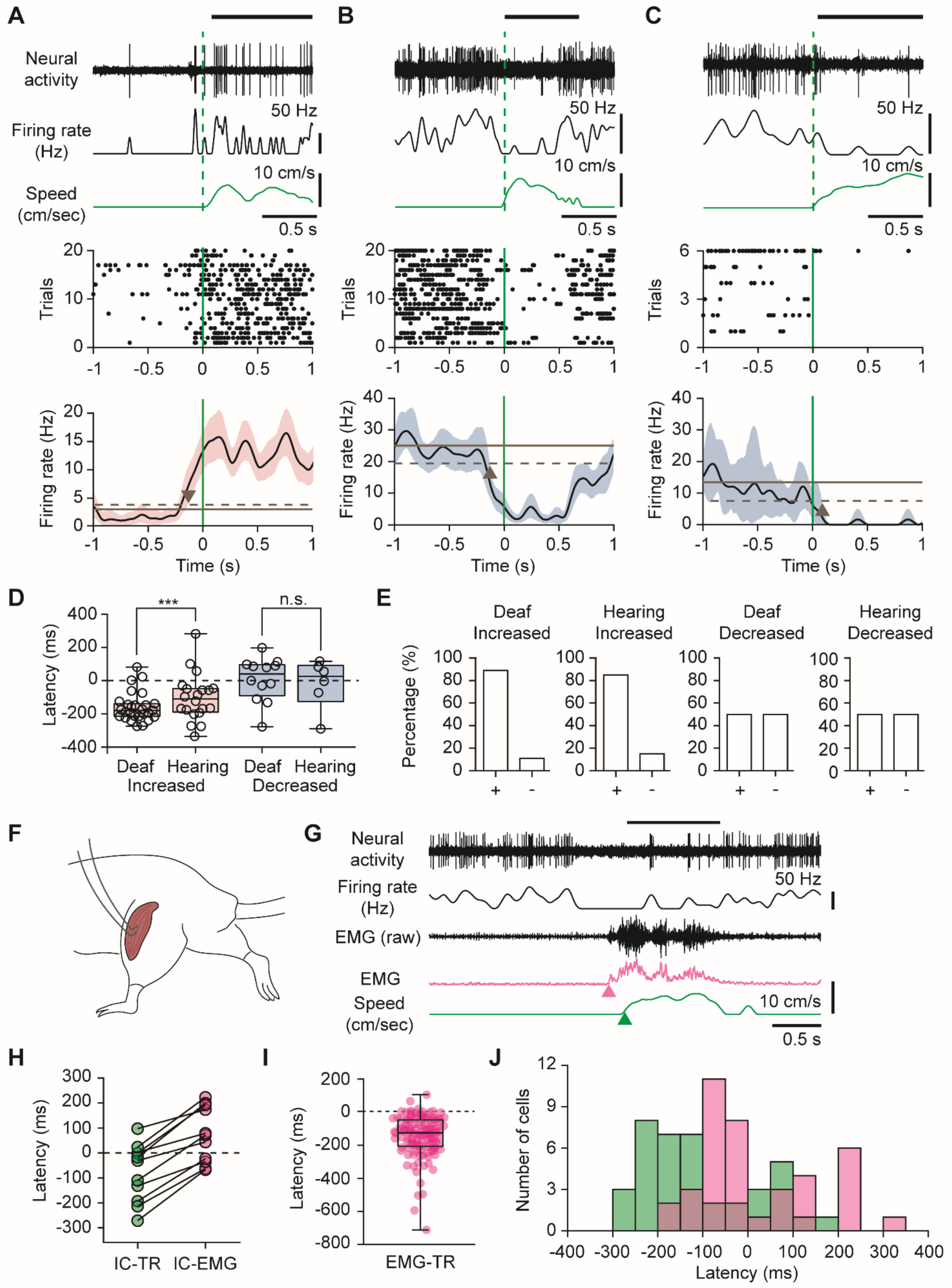
Modulation latency of locomotion-related IC activity in deaf mice. (**A-C**) Examples of IC neurons showing increased (A) or decreased firing (B, C) at locomotion onset in deaf mice. Top panels: Raw neural activity (top trace), smoothed firing rate (middle trace), and walking speed (bottom trace, green). Black horizontal bars, above the top trace, indicate walking periods; green dashed vertical line marks locomotion onset. Middle panels: Raster plots of spike times aligned to locomotion onset across bouts, with green vertical line marking the locomotion onset. Bottom panels: Locomotion onset-triggered average firing rates (black traces) with the 95% confidence intervals (pink or blue shaded area). Horizontal solid and dashed lines indicate the mean ± 2 SD of baseline activity. Arrowheads indicate the latency of modulation, defined as the time at which the confidence interval first crosses the 2 SD threshold. (**D**) Comparison of modulation latencies between deaf and hearing mice for neurons with increased (left; Deaf: n = 27 neurons; Hearing: n = 20 neurons) and decreased (right; Deaf: n = 12 neurons; Hearing: n = 6 neurons) activity. *** p<0.001, n.s., not significant (Mann-Whitney U test). (**E**) Proportion of neurons with modulation preceding (+) or following (-) locomotion onset in deaf and hearing mice. (**F**) Schematic of EMG recording for the hindlimb muscle activity. (**G**) Simultaneous recording of IC activity, EMG, and treadmill speed during a walking bout. From top to bottom: raw neural activity, smoothed firing rate, raw and smoothed EMG signal (magenta), and walking speed (green). (**H**) Latency of IC neural modulation relative to treadmill onset (IC-TR) and EMG onset (IC-EMG) for neurons recorded with EMG (n = 11). (**I**) Delays between EMG and treadmill onsets across bouts (n = 121 waking bouts from 11 mice). Box and whiskers plot with min.& max. data are shown. (**J**) Distribution of IC modulation latencies aligned to treadmill (green) and EMG (magenta) onsets.

Due to the delay between limb muscle activation and the actual movement onset of the treadmill, some of the modulation signals that appear to precede locomotion onset may, in fact, follow muscle activation. To obtain more accurate timing measurements, we also recorded electromyographic (EMG) signals from the hindlimb muscles during a subset of IC recordings (Fig 3F; see Methods).

As illustrated by the example of a suppressed neuron, the onset of hindlimb EMG signals preceded treadmill movement onset (Fig 3G). In a subset of IC neurons in which both treadmill and EMG signals were recorded (n = 11 neurons; 5 with increased and 6 with decreased activity), the median modulation latency shifted by 102 ms, from −30 ms relative to treadmill onset to 72 ms relative to EMG onset (Fig 3H). To better estimate modulation latency relative to EMG onset, we measured the time difference between EMG and treadmill onsets across many movement bouts. This difference was variable, ranging from −593 ms to 6 ms, with a median value of −125 ms (Fig 3I; n = 121 bouts from 11 mice). Applying this correction to all neurons shifted the median modulation latency from −147 ms (treadmill based) to −22 ms (EMG based; Fig 3J). This shift toward positive latency values increased the proportion of neurons with positive latency values from 23% to 39% (Fig 3J), suggesting that a larger portion is consistent with feedback signals than previously suggested.

Each IC neuron may receive a unique combination of predictive and feedback-related signals, resulting in heterogeneous modulation patterns. Supporting this idea, when neural activity was aligned at locomotion onset, we observed substantial variability in both the latency and direction of modulation across different walking bouts, even within the same neuron (Fig 4A and 4B). In neurons where bout-wise analysis was feasible (see Methods), modulation onset varied substantially, often straddling the movement onset (median range for increased neurons: 327 ms (n = 7); decreased neurons: 306 ms (n = 4)) (Fig 4C and 4D). We also asked whether the direction of modulation varied within individual neurons; i.e., if suppression occurs in a neuron with increased average firing rate or vice versa. In 6 of 11 neurons, we observed bouts with modulation in the opposite direction to the neuron’s average change, occurring in ∼14 ± 5% of bouts (Fig 4E). These findings support the idea that individual IC neurons integrate diverse set of inputs during locomotion.

**Fig 4.**
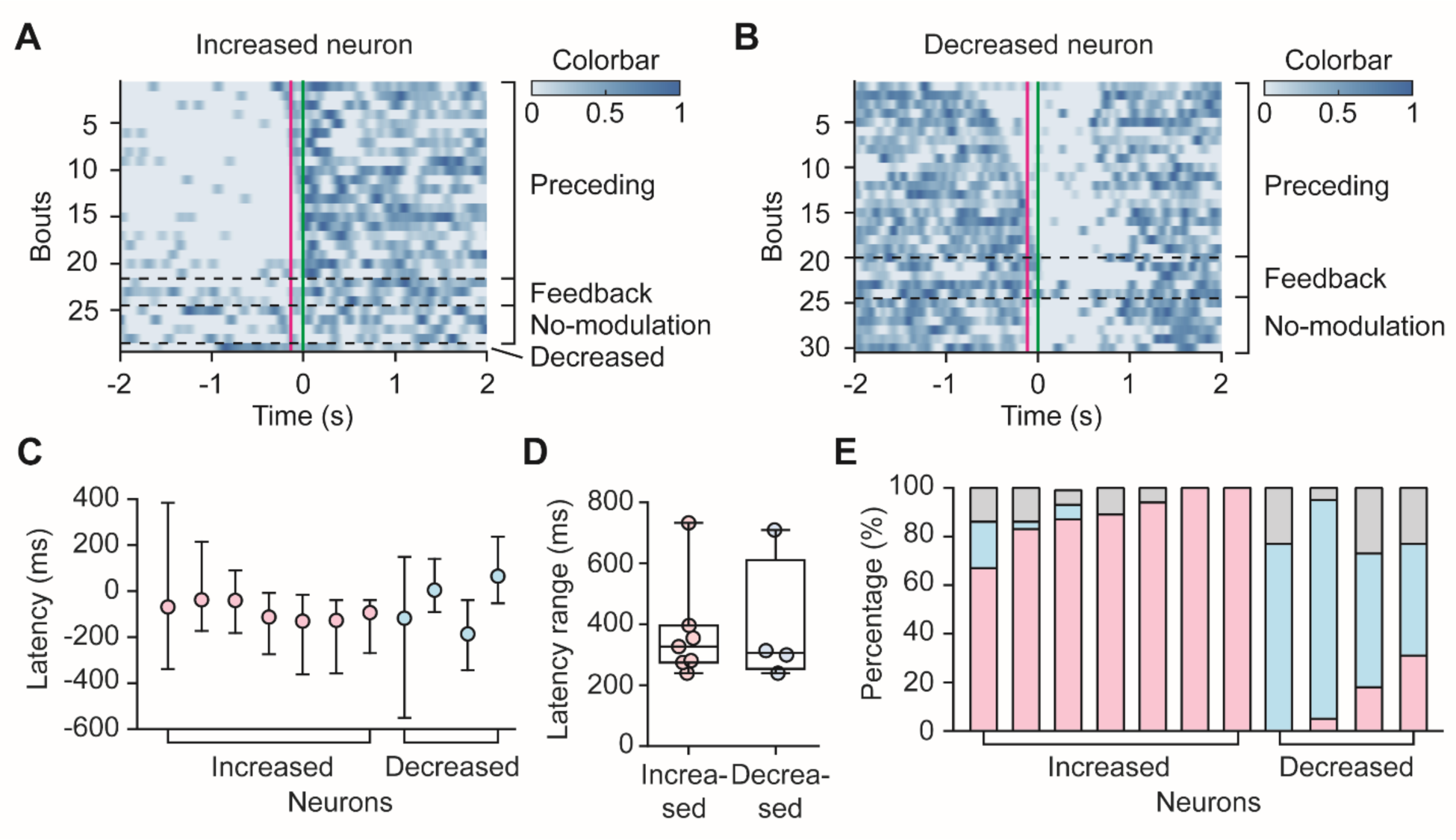
Variability in IC neural modulation across locomotion bouts. (**A**, **B**) Heatmaps of normalized firing rates from individual locomotion bouts for example neurons showing overall increased (A) or decreased (B) firing rates during walking. Each row represents one locomotion bout, aligned to locomotion onset (green vertical line). Movement onset after applying the median EMG-treadmill delay (Fig 3I) is also shown (magenta vertical line). Bouts are sorted and grouped based on modulation type (Preceding, Feedback, No-modulation, Decreased), with black dashed lines indicating group boundaries. (**C**) Mean modulation latency for each neuron with overall increased (left, *n* = 7) or decreased (right, *n* = 4) firing during walking. Error bars indicate the range across bouts. (**D**) Range of modulation latencies across bouts for the neurons in (C). (**E**) Proportion of bout-wise modulation types for each neuron. Stacked bars show the percentage of bouts exhibiting increased (pink), decreased (blue), or no modulation (gray) in firing relative to locomotion onset.

## DISCUSSION

To understand the mechanisms underlying locomotion-induced neural modulation in the IC, it is crucial to isolate the contributions of distinct input sources. In this study, using a mouse line that can be rendered deaf, we were able to separate non-auditory, movement-related neural signals from both air- and bone-conducted auditory feedback. We found that this modulation persists even in the absence of auditory input, with both excitatory and suppressive changes observed across all IC subdivisions. By defining locomotion onset using hindlimb EMG, rather than treadmill motion, we were able to more precisely establish that non-auditory modulation includes both predictive and feedback components. Additionally, we observed substantial variability in modulation timing and direction across locomotion bouts, consistent with the involvement of diverse non-auditory sources. The widespread distribution of this modulation, including within the CNIC, supports the idea that integrating non-auditory, movement-related information with auditory input is a core feature of IC function.

### Isolation of non-auditory neural modulation in the IC

Our results in deaf mice provide clear evidence that IC neurons exhibit robust non-auditory modulation during locomotion. By comparing modulation patterns between hearing and deaf mice under comparable conditions, we estimate that approximately 20% of excitatory modulation observed in hearing mice can be attributed to auditory feedback – an effect not separable in previous studies using hearing mice [6,7]. In contrast, the proportion of neurons exhibiting suppressive modulation remained unchanged in deaf mice, suggesting that the suppression likely is driven by non-auditory inputs. Moreover, the persistence of modulation in the CNIC of deaf animals suggests that modulation in this region in hearing mice is not solely due to auditory feedback. The contribution of air-conducted auditory feedback may vary with environmental conditions, and it would be important to investigate how non-auditory signals interact with auditory feedback under different behavioral and sensory contexts [22,30].

If locomotion generates skull vibrations, these could propagate through the auditory pathway via bone conduction. Indeed, we detected consistent movement-related skull vibrations even under head-fixed conditions, revealing a previously underappreciated source of auditory feedback. In deaf mice, this bone-conducted component is also eliminated, allowing a clear separation of non-auditory modulation. Although the precise contribution of bone conduction remains difficult to quantify, it is important to recognize that such signals would persist in the presence of masking noise – an approach commonly used to investigate movement-related, non-auditory neural activity [1,31].

Our recordings were performed approximately one week after deafening, raising the question of whether neural plasticity could influence the results. While adult-onset hearing loss is known to induce plastic changes in auditory and multisensory brain regions [32,33], these changes tend to involve the reorganization of existing circuits rather than the formation of entirely new connections. Furthermore, recent work has shown that motor-related signals in the auditory cortex are preserved in both congenitally deaf and hearing mice, suggesting that such signals are not highly dependent on auditory experience [34]. While we cannot completely rule out plastic changes in movement-related IC inputs following deafening, these are likely to be quantitative rather than qualitative in nature.

### Predictive vs feedback signals

The timing of neural modulation relative to locomotion onset offers insights into whether a signal is predictive or feedback-driven. Typically, movement onset is defined by treadmill motion; however, we show that limb muscle activation (as measured by EMG) precedes treadmill motion by approximately 150 ms. Applying this correction shifts the estimated onset of neural modulation closer to muscle activation (from −147 ms to −22 ms) and reduces the fraction of signals classified as predictive (Fig. 3). Even with this adjustment, many IC neurons exhibit modulation that clearly precedes movement onset, indicating the presence of predictive signals. Potential source of such inputs include midbrain locomotion circuits [35], motor cortex [36,37], and neuromodulatory systems [38]. We also found evidence for non-auditory feedback signals that follows limb muscle activation. Conversely, delayed modulation relative to EMG likely reflects somatosensory or proprioceptive feedback signals activated during movement [28,29,39–41]. Once locomotion begins, predictive and feedback signals overlap temporally, making them difficult to disentangle. Dissecting these overlapping sources will require targeted manipulation of input pathways, which would be aided by the ability to exclude auditory components, as in our current approach.

### Movement and the auditory pathway

Our findings raise broader questions about how movement information is integrated throughout the auditory pathway to support behavior. Neural activity in auditory cortical neurons is known to be modulated by locomotion and other motor behaviors [42]. While the sources of such modulation may differ depending on specific behavior, accumulating evidence points to direct non-auditory inputs to the auditory cortex, rather than inheritance from the auditory thalamus [1,31,38]. On the other hand, recent evidence including ours results demonstrate that movement-related modulation is already prominent in subcortical auditory regions [4,6,7,43]. The relationship between the subcortical and cortical modulations remains to be elucidated. It is possible that different stages of the auditory pathway may encode movement via distinct mechanisms. For example, the IC receives direct ascending and descending somatosensory inputs, which are likely activated during locomotion [29,40]. In addition, MGB does not mirror IC modulation patterns: locomotion primarily attenuates sound-evoked responses in the medial geniculate body (MGB) without substantially affecting baseline activity [4], whereas the IC exhibits robust modulation in both spontaneous and evoked firing. The anatomical and physiological differences raise the possibility that locomotion-induced modulations arise from different mechanisms and may even serve heterogeneous functions across auditory processing stages. The functional role of movement modulation in the auditory pathway remains controversial. Recent studies suggest that auditory cortical neurons encode running speed and prediction error, indicating a role in monitoring self-generated movements and their sensory consequences [10,30]. Our findings suggest that to some degree, such encoding already occurs at the midbrain level. This aligns with emerging evidence that IC neurons are responsive to a broad range of behavioral variables beyond sound, including attention, arousal, and eye movement [44–46]. Tracking movement information at early stages of auditory processing may allow for rapid responses to sounds encountered during navigation, supporting adaptive behavior in dynamic and complex environments.

## MATERIALS AND METHODS

**Resources table**

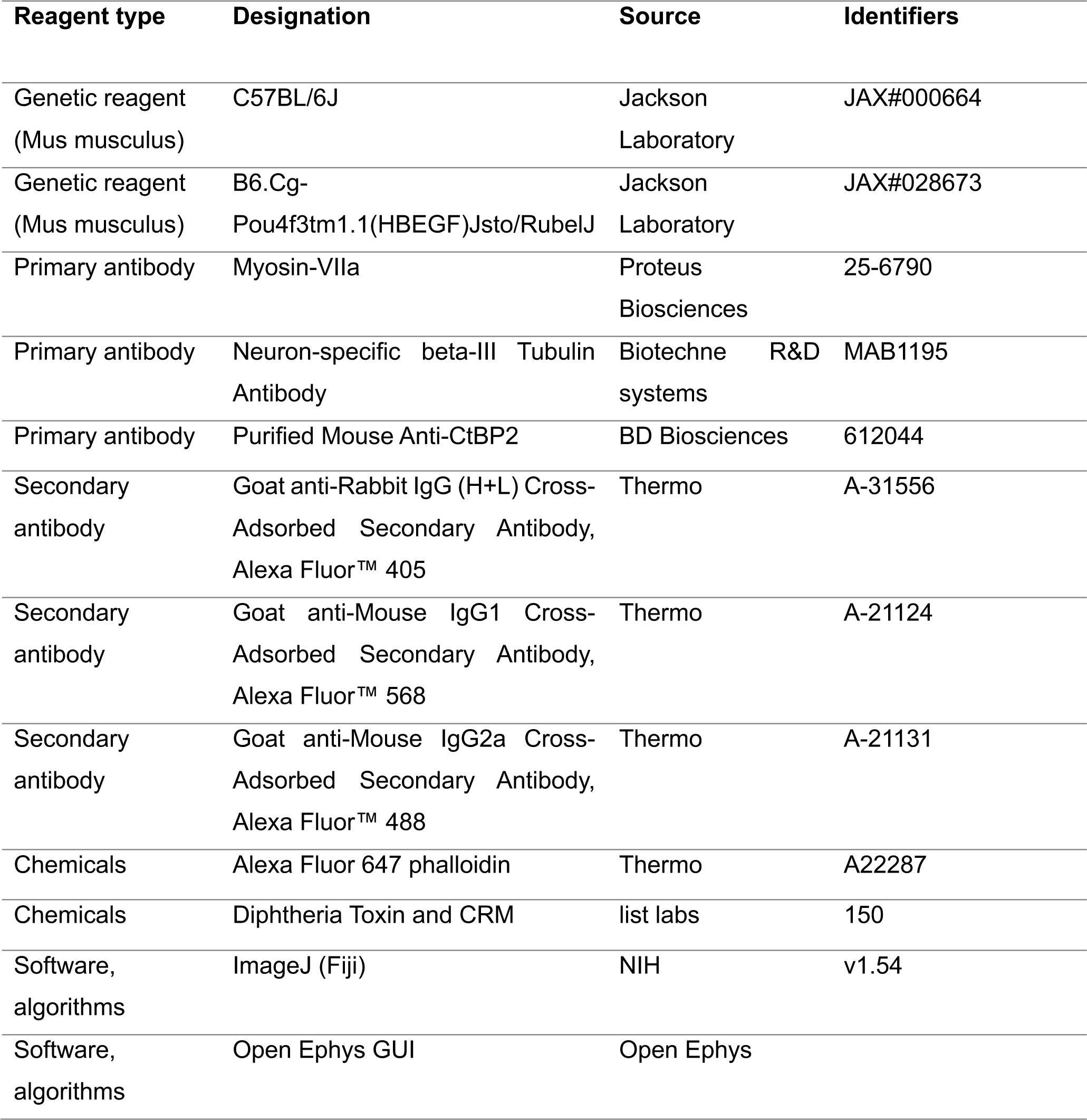

### Animal preparation

All procedures were approved by the Institutional Animal Care and Use Committee (IACUC) of the Korea Brain Research Institute (IACUC-23-00077-M2) and complied with institutional guidelines. Mice were housed under a 12:12-hour light/dark cycle with ad libitum access to food and water. Experiments were conducted using C57BL/6J mice (N = 16) and Pou4f3^+/DTR^ mice (N = 19) of both sexes, aged 6-15 weeks. Deafness was induced in Pou4f3^+/DTR^ mice via a single intramuscular injection of diphtheria toxin (25 ng/kg) at least 7 days prior to neural recording.

### Headpost surgery

Mice were anesthetized via intraperitoneal injection of ketamine (0.10 mg/g, Yuhan) and xylazine (0.01 mg/g, Bayer Korea). After shaving the scalp, the head was secured in a stereotaxic apparatus, and lidocaine was injected subcutaneously for local anesthesia. A midline incision was made to expose the skull, and the periosteum was removed. The headpost was positioned over the IC region (1.0 mm lateral, 5.0 mm posterior to bregma) and secured with dental cement.

### Skull vibration measurement

Skull vibrations were recorded using a vibration sensor (BU-21771-000, Knowles, IL, USA), affixed to the exposed skull with UV-cured dental adhesive. The accelerometer was powered by a 1.6 V AA battery, and signals were acquried using an Intan headstage (RHD2132, Intan Technologies) and the Open Ephys acquisition system. The raw signal (5 Hz–16 kHz detection range) was bandpass filtered (20–450 Hz) and filtered to remove 60 Hz noise. To extract the envelope, the signal was downsampled from 30,000 Hz to 1,000 Hz, rectified, and smoothed using a moving average filter.

### Cochlear histology

Following transcardial perfusion with 4% paraformaldehyde, cochleae were extracted, post-fixed overnight at 4°C, and decalcified in 125 mM EDTA for 3–5 days. The cochlea was separated into basal, middle, and apical turns. Samples were blocked for 2 hours at room temperature and incubated with primary antibodies overnight at 4°C. The following antibodies were used: CtBP2 (1:2000, mouse) for hair cell presynaptic ribbons, Tuj1 (1:2000, mouse) for afferent fibers, and Myo7a (1:1000, rabbit) for hair cells. Alexa Fluor-conjugated secondary antibodies (1:1000) were used, and Alexa Fluor 405-Phalloidin (1:50) was used for F-actin labeling. Sections were imaged using a Leica TCS SP8 confocal microscope (40× objective) and processed in ImageJ.

### IC neural recordings

Extracellular recordings were conducted in awake, head-fixed mice after at least 7 days of treadmill training (30 min/day). A craniotomy (∼3 mm diameter) over the IC (1.0 mm lateral, 5.0 mm posterior to bregma) was performed under isoflurane anesthesia. After recovery, a tungsten electrode array (∼10 MΩ, 6 electrodes with ∼200 µm spacing, FHC) was inserted at 100–200 µm posterior to transverse sinus and 0.4–2.0 mm lateral to the midline, reaching depths of up to 1800 µm. Neural activity was recorded using a 16-channel headstage (RHD2132, Intan Technologies) and the Open Ephys acquisition system at 30 kHz and bandpass filtered between 600–6000 Hz. Treadmill signal was simultaneously recorded using an optical encoder (Scitech Korea, Seoul, Korea).

### Data analysis

To confirm deafness, white noise bursts (2–64 kHz, 10–90 dB SPL, 50 ms duration) were presented while mapping the IC with a multi-electrode array. Multi-unit spikes exceeding 3× baseline standard deviation were extracted, and sound-evoked responses were quantified as average spike rates during noise stimulus presentation (Fig 2B, 2C and 1S Fig).

For IC recordings during locomotion, single units were isolated using Offline Sorter v4 (Plexon). Spike waveforms were clustered via principal component analysis (PCA), and only clearly separable clusters were accepted as single units (multivariate ANOVA with *p* < 0.05). Only units with <0.1% refractory period violations (inter-spike interval < 0.7 ms) were classified as single neurons.

Spontaneous activity was analyzed from ≥200 s of silent recordings, divided into 1-second segments and categorized as stationary (0 cm/s), walking (>2 cm/s), or transitional. Only purely stationary or walking segments were included. Neural modulation was defined as a significant difference (p < 0.01) in firing rate between walking and stationary periods across at least 4 walking segments. Walking speed was derived by smoothing the treadmill signal using a 200 ms Hanning filter. Modulation latency was determined by aligning neural activity to locomotion onset, defined as a >5% change in treadmill signal following ≥0.5 s of stable baseline. For each bout, the smoothed firing rate and 95% confidence interval were estimated via bootstrap resampling. Latency was defined as the time at which lower or upper bound of confidence interval deviated by more than ±2 SD from baseline, relative to the onset. Only neurons with ≥4 qualified bouts were included. For bout-wise latency analysis (Fig 4), each bout was required to have ≥0.5 s of stable baseline before locomotion onset.

### Histology for recording locations

At the conclusion of neural recordings, electrolytic lesions were made at depths of 500 µm and 1000 µm (30 µA, 10 s x 2). Mice were perfused, and brains were extracted. Brains were post-fixed, cryoprotected in 30% sucrose, embedded in OCT, and sectioned at 40 µm using a cryostat. Nissl staining was performed, and sections were imaged using a slide scanner (Pannoramic Scan). Recording sites were verified based on the locations of lesions in Nissl images (Fig 2I).

### Electromyography (EMG) recording

EMG signals were recorded from the right hindlimb using PFA-coated stainless steel wire (bare diameter: 0.003“, Cat. No. 791100, A-M Systems) or tungsten wire (bare diameter: 0.005”, Cat. No. 796500, A-M Systems). A ∼3 mm hooked wire tip was inserted into the muscle through a syringe needle. Signals were acquired using an Intan headstage (RHD2132, Intan Technologies) and the Open Ephys system. Data were bandpass filtered (20–450 Hz), notch filtered at 60Hz (±5 Hz), and downsampled to 2,000 Hz. To obtain the envelope, EMG traces were rectified and smoothed using a moving average filter. EMG onset was determined when the smoothed EMG signal exceeded the mean plus three standard deviations of the baseline activity measured over a 0.5–1 second period.

### Statistical analysis

All statistical analyses were performed in MATLAB. Unless otherwise indicated, statistical significance was defined as p < 0.05. For comparisons of spontaneous activity between stationary and walking periods, a threshold of p < 0.01 (Student’s t-test) was used. Welch’s t-test was used for comparing group means with unequal sample sizes (Fig 2C, S2A and S2C Figs). Paired t-tests were used for within-subject comparisons (e.g., walking vs. stationary; S2B Fig). For comparisons involving medians or non-normally distributed small-sample data, Mann–Whitney U-tests were used (S2C Fig, Fig. 3D).

## ACKNOWLEDGEMENT

This project was funded by Korea Brain Research Institute (25-BR-01-01 and 25-BR-07-01), and National Research Foundation of Korea (NRF-2021R1F1A1049434).

## AUTHOR CONTRIBUTIONS

Author contributions

---

**S1 Fig.**
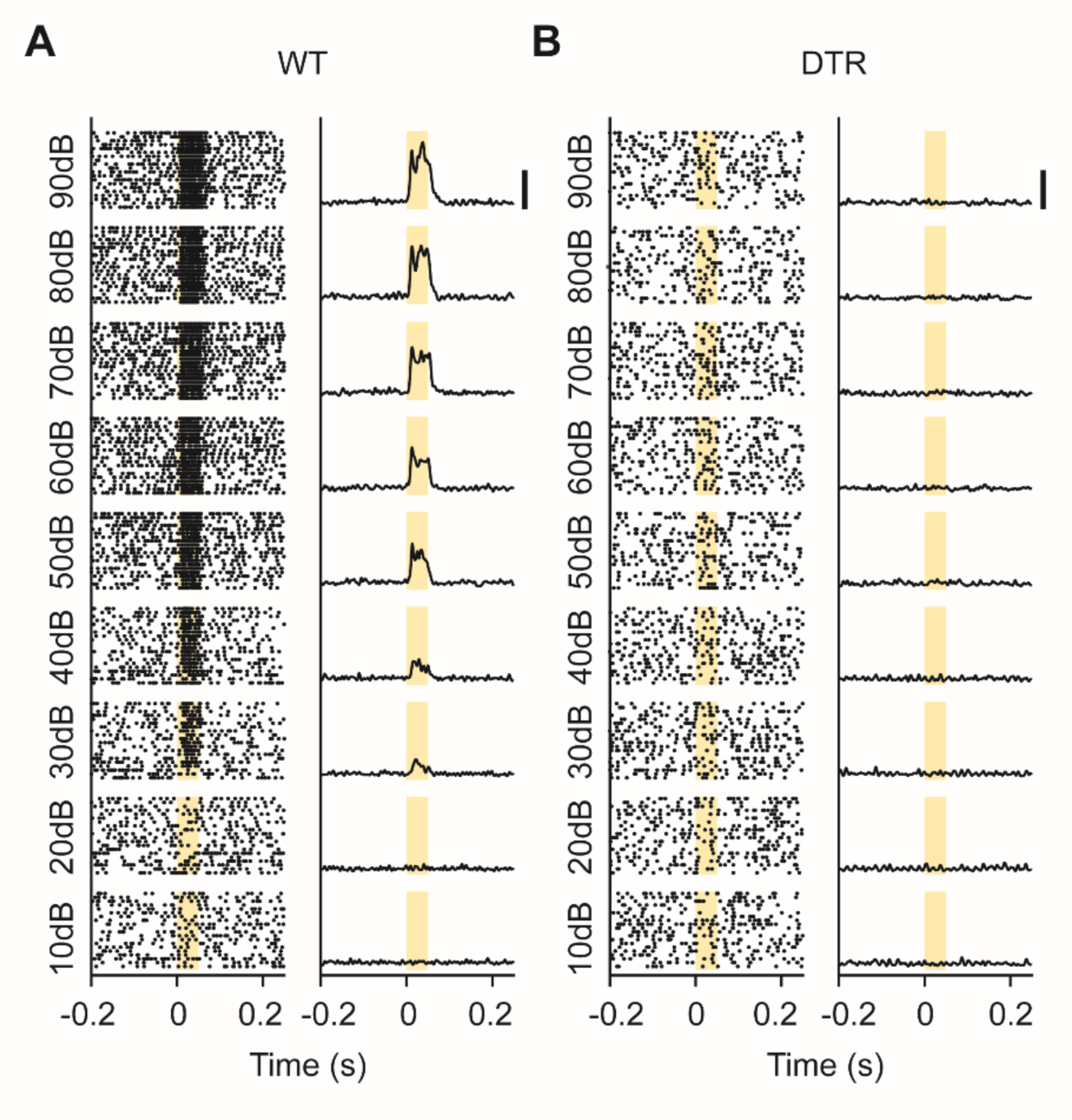
Verification of deafness in Pou4f3^+/DTR^ mice following DT injection. (**A, B**) Representative multi-unit recordings from IC neurons in response to a broadband sound (50 ms, 10-90 dB SPL) in a WT mouse (A) and a DTR mouse following DT injection (B). Left panels: Raster plots showing spike times across repeated trials for each stimulus intensity. Right panels: Corresponding peristimulus time histograms (PSTHs) showing average firing rates over time. Yellow shades indicate the duration of the sound stimulus. The multi-unit data are from one of the recording sites shown in Fig 2B. Robust sound-evoked responses are observed in WT mice, but are absent in DTR mice (Fig 2B and 2C). Scale bar, 500 Hz.

**S2 Fig.**
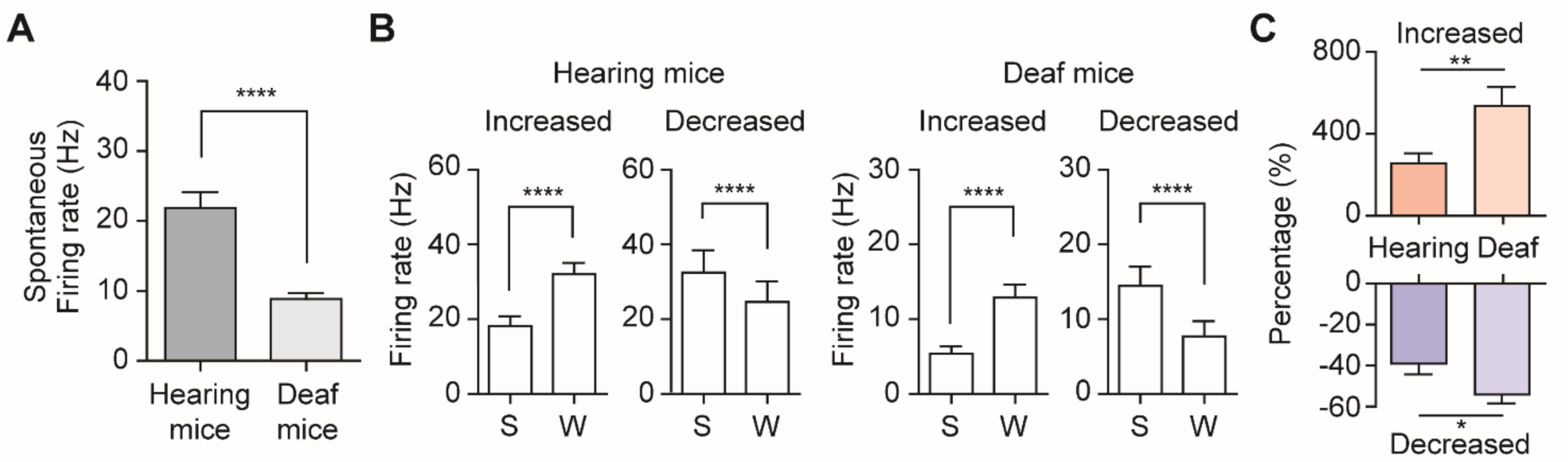
Modulation of IC neural activity during locomotion in hearing and deaf mice. (**A**) Average spontaneous firing rates in hearing (n = 96 neurons) and deaf mice (n = 146 neurons). Deaf mice exhibited significantly lower baseline firing rates (hearing: 21.9 ± 2.2 Hz; deaf: 8.8 ± 0.9 Hz; mean ± SEM; ****p < 0.0001, unpaired t-test with Welch’s correction). (**B**) Average firing rates during stationary (S) and walking (W) periods in neurons with increased or decreased firing. Locomotion significantly modulated firing in both hearing (left; n = 51 increased, n = 22 decreased) and deaf (right; n = 64 increased, n = 32 decreased) mice (error bars, SEM; **** p < 0.0001, paired t-test). (**C**) Percent change in firing rate from stationary to walking periods for neurons with increased (top) or decreased firing (bottom). Modulation was significantly greater in deaf mice (Increased: 257% vs 537%, ** p = 0.0081, Unpaired t test with Welch’s correction; Decreased: −39% vs −54%, * p = 0.0394, Mann-Whitney test). Error bars denote SEM.

**S3 Fig.**
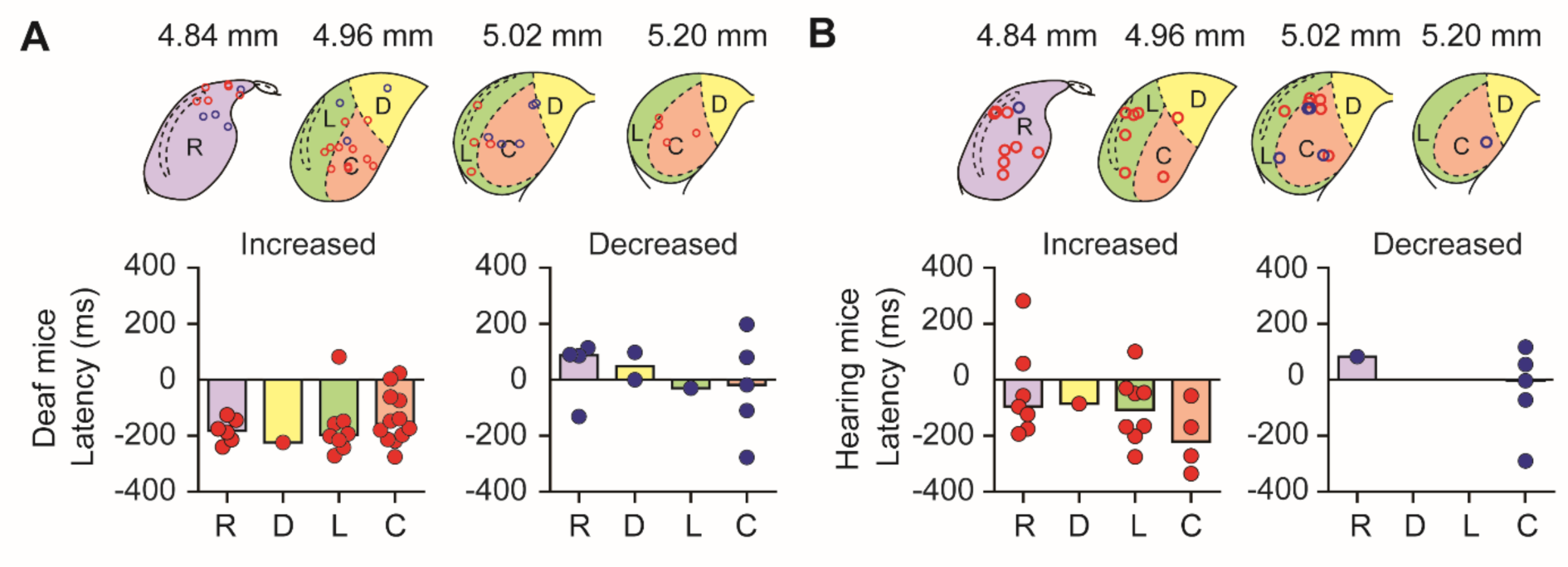
Spatial distribution of locomotion-induced modulation latencies in the IC of deaf and hearing mice. (**A**, **B**) Latency of locomotion-induced modulation in IC neurons, separated by IC subregions in deaf (A) and hearing (B) mice. Top: Schematic diagrams of IC subdivisions (R: rostral, D: dorsal, L: lateral, C: caudal) at four rostro-caudal positions. Red and blue circles represent neurons with increased and decreased firing rates during locomotion, respectively. Bottom: Latency distributions of modulated neurons within each subregion. Colored bars reflect mean latencies for each sub-region. (**A**) In deaf mice, median latencies of increased neurons were: rostral −182 ms, dorsal −224 ms, lateral −197 ms, and caudal −160 ms. Decreased neurons exhibited median latencies of: rostral 88 ms, dorsal 49 ms, lateral −30 ms, and caudal −18 ms. (**B**) In hearing mice, increased neurons had median latencies of: rostral −96 ms, dorsal −85 ms, lateral −107 ms, and caudal −221 ms. Decreased neurons showed latencies of: rostral 83 ms and caudal −3 ms.

## REFERENCES

1. Schneider DM, Nelson A, Mooney R. A synaptic and circuit basis for corollary discharge in the auditory cortex. Nature. 2014;513: 189–194. doi:10.1038/nature13724

2. Zhou M, Liang F, Xiong XR, Li L, Li H, Xiao Z, et al. Scaling down of balanced excitation and inhibition by active behavioral states in auditory cortex. Nat Neurosci. 2014;17: 841–850. doi:10.1038/nn.3701

3. McGinley MJ, Vinck M, Reimer J, Batista-Brito R, Zagha E, Cadwell CR, et al. Waking State: Rapid Variations Modulate Neural and Behavioral Responses. Neuron. Cell Press; 2015. pp. 1143–1161. doi:10.1016/j.neuron.2015.09.012

4. Williamson RS, Hancock KE, Shinn-Cunningham BG, Polley DB. Locomotion and Task Demands Differentially Modulate Thalamic Audiovisual Processing during Active Search. Current Biology. 2015;25: 1885–1891. doi:10.1016/j.cub.2015.05.045

5. Bigelow J, Morrill RJ, Dekloe J, Hasenstaub AR. Movement and VIP interneuron activation differentially modulate encoding in mouse auditory cortex. eNeuro. 2019;6. doi:10.1523/ENEURO.0164-19.2019

6. Chen C, Song S. Differential cell-type dependent brain state modulations of sensory representations in the non-lemniscal mouse inferior colliculus. Commun Biol. 2019;2. doi:10.1038/s42003-019-0602-4

7. Yang Y, Lee J, Kim G. Integration of locomotion and auditory signals in the mouse inferior colliculus. Elife. 2020;9. doi:10.7554/eLife.52228

8. Henschke JU, Price AT, Pakan JMP. Enhanced modulation of cell-type specific neuronal responses in mouse dorsal auditory field during locomotion. Cell Calcium. 2021;96. doi:10.1016/j.ceca.2021.102390

9. Khoury CF, Fala NG, Runyan CA. Arousal and Locomotion Differently Modulate Activity of Somatostatin Neurons across Cortex. eNeuro. 2023;10. doi:10.1523/ENEURO.0136-23.2023

10. Vivaldo CA, Lee J, Shorkey MC, Keerthy A, Rothschild G. Auditory cortex ensembles jointly encode sound and locomotion speed to support sound perception during movement. PLoS Biol. 2023;21. doi:10.1371/journal.pbio.3002277

11. Morandell K, Yin A, Rio RT Del, Schneider DM. Movement-Related Modulation in Mouse Auditory Cortex Is Widespread Yet Locally Diverse. Journal of Neuroscience. 2024;44. doi:10.1523/JNEUROSCI.1227-23.2024

12. Rummell BP, Klee JL, Sigurdsson T. Attenuation of responses to self-generated sounds in auditory cortical neurons. Journal of Neuroscience. 2016;36: 12010–12026. doi:10.1523/JNEUROSCI.1564-16.2016

13. Schneider DM, Sundararajan J, Mooney R. A cortical filter that learns to suppress the acoustic consequences of movement. Nature. 2018;561: 391–395. doi:10.1038/s41586-018-0520-5

14. Audette NJ, Zhou WX, La Chioma A, Schneider DM. Precise movement-based predictions in the mouse auditory cortex. Current Biology. 2022;32: 4925–4940.e6. doi:10.1016/j.cub.2022.09.064

15. Malmierca MS. The inferior colliculus: A center for convergence of ascending and descending auditory information. Neuroembryology and Aging. 2004. pp. 215–229. doi:10.1159/000096799

16. Jeffery A. Winer, Christoph E. Schreiner. The Inferior Colliculus. Springer; 2005.

17. Aitkin LM, Dickhaus H, Schult W, Zimmermann M. External Nucleus of Inferior Colliculus: Auditory and Spinal Somatosensory Merents and Their Interactions. JOURNALOF NEUROPHYSIOLOGY. 1978.

18. Aitkin LM, Kenyon CE, Philpcyl’t P. The Representation of the Auditory and Somatosensory Systems in the External Nucleus of the Cat Inferior Colliculus. J Comp Neurol. 1981.

19. Morest KD, Oliver DL. The Neuronal Architecture of the Inferior Colliculus in the Cat: Defining the Functional Anatomy of the Auditory Midbrain. Journal of Comparative Neurology. 1984; 222– 209.

20. Coleman JR, Cleric1 WJ. Sources of Projections to Subdivisions of the Inferior Colliculus in the Rat. Journal of Comparative Neurology. 1987;262: 215–226.

21. Lesicko AMH, Hristova TS, Maigler KC, Llano DA. Connectional modularity of top-down and bottom-up multimodal inputs to the lateral cortex of the mouse inferior colliculus. Journal of Neuroscience. 2016;36: 11037–11050. doi:10.1523/JNEUROSCI.4134-15.2016

22. Cornwell T, Woodward J, Wu M, Jackson B, Souza P, Siegel J, et al. Walking With Ears: Altered Auditory Feedback Impacts Gait Step Length in Older Adults. Front Sports Act Living. 2020;2. doi:10.3389/fspor.2020.00038

23. Puria S, Rosowski JJ. Békésy’s contributions to our present understanding of sound conduction to the inner ear. Hearing Research. 2012. pp. 21–30. doi:10.1016/j.heares.2012.05.004

24. Tong L, Strong MK, Kaur T, Juiz JM, Oesterle EC, Hume C, et al. Selective deletion of cochlear hair cells causes rapid age-dependent changes in spiral ganglion and cochlear nucleus neurons. Journal of Neuroscience. 2015;35: 7878–7891. doi:10.1523/JNEUROSCI.2179-14.2015

25. Fukushima M, Margoliash D. The effects of delayed auditory feedback revealed by bone conduction microphone in adult zebra finches. Sci Rep. 2015;5. doi:10.1038/srep08800

26. Fettiplace R. Hair cell transduction, tuning, and synaptic transmission in the mammalian cochlea. Compr Physiol. 2017;7: 1197–1227. doi:10.1002/cphy.c160049

27. Malmierca MS, Blackstad TW, Osen KK. Computer-assisted 3-D reconstructions of Golgi-impregnated neurons in the cortical regions of the inferior colliculus of rat. Hear Res. 2011;274: 13–26. doi:10.1016/j.heares.2010.06.011

28. Akay T, Tourtellotte WG, Arber S, Jessell TM. Degradation of mouse locomotor pattern in the absence of proprioceptive sensory feedback. Proc Natl Acad Sci U S A. 2014;111: 16877–16882. doi:10.1073/pnas.1419045111

29. Turecek J, Ginty DD. Coding of self and environment by Pacinian neurons in freely moving animals. Neuron. 2024. doi:10.1016/j.neuron.2024.07.008

30. Solyga M, Keller GB. Multimodal mismatch responses in mouse auditory cortex. eLife. 2025. doi:10.7554/eLife.95398.3

31. Clayton KK, Williamson RS, Hancock KE, Tasaka G ichi, Mizrahi A, Hackett TA, et al. Auditory Corticothalamic Neurons Are Recruited by Motor Preparatory Inputs. Current Biology. 2021;31: 310–321.e5. doi:10.1016/j.cub.2020.10.027

32. Salvi RJ, Wang J, Ding D. Auditory plasticity and hyperactivity following cochlear damage. Hear Res. 2000;147: 261–274. Available: www.elsevier.com/locate/heares

33. Schormans AL, Scott KE, Vo AMQ, Tyker A, Typlt M, Stolzberg D, et al. Audiovisual temporal processing and synchrony perception in the rat. Front Behav Neurosci. 2017;10. doi:10.3389/fnbeh.2016.00246

34. Harmon TC, Madlon-Kay S, Pearson J, Mooney R. Vocalization modulates the mouse auditory cortex even in the absence of hearing. Cell Rep. 2024;43. doi:10.1016/j.celrep.2024.114611

35. Lee AM, Hoy JL, Bonci A, Wilbrecht L, Stryker MP, Niell CM. Identification of a brainstem circuit regulating visual cortical state in parallel with locomotion. Neuron. 2014;83: 455–466. doi:10.1016/j.neuron.2014.06.031

36. Olthof BMJ, Rees A, Gartside SE. Multiple nonauditory cortical regions innervate the auditory midbrain. Journal of Neuroscience. 2019;39: 8916–8928. doi:10.1523/JNEUROSCI.1436-19.2019

37. Gartside SE, Olthof BM, Rees A. Motor, somatosensory, and executive cortical areas elicit monosynaptic and polysynaptic neuronal activity in the auditory midbrain. Hear Res. 2024;447. doi:10.1016/j.heares.2024.109009

38. Reimer J, McGinley MJ, Liu Y, Rodenkirch C, Wang Q, McCormick DA, et al. Pupil fluctuations track rapid changes in adrenergic and cholinergic activity in cortex. Nat Commun. 2016;7. doi:10.1038/ncomms13289

39. Huey EL, Turecek J, Delisle MM, Mazor O, Romero GE, Dua M, et al. The auditory midbrain mediates tactile vibration sensing. Cell. 2025;188: 104–120.e18. doi:10.1016/j.cell.2024.11.014

40. Lohse M, Dahmen JC, Bajo VM, King AJ. Subcortical circuits mediate communication between primary sensory cortical areas in mice. Nat Commun. 2021;12. doi:10.1038/s41467-021-24200-x

41. Lohse M. Integration of somatosensory and motor-related information in the auditory system. Frontiers in Neuroscience. Frontiers Media S.A.; 2022. doi:10.3389/fnins.2022.1010211

42. Schneider DM, Mooney R. How Movement Modulates Hearing. 2018. doi:10.1146/annurev-neuro-072116

43. Singla S, Dempsey C, Warren R, Enikolopov AG, Sawtell NB. A cerebellum-like circuit in the auditory system cancels responses to self-generated sounds. Nat Neurosci. 2017;20: 943–950. doi:10.1038/nn.4567

44. Lee T-Y, Weissenberger Y, King AJ, Dahmen JC. Midbrain encodes sound detection behavior without auditory cortex. Elife. 2024;12. doi:10.7554/eLife.89950

45. Du X, Xu H, Song P, Zhai Y, Ye H, Bao X, et al. Beyond Auditory Relay: Dissecting the Inferior Colliculus’s Role in Sensory Prediction, Reward Prediction and Cognitive Decision-Making. 2025. doi:10.7554/eLife.101142.2

46. Gruters KG, Groh JM. Sounds and beyond: Multisensory and other non-auditory signals in the inferior colliculus. Frontiers in Neural Circuits. 2012. pp. 1–39. doi:10.3389/fncir.2012.00096

